# Growth-Altering Microbial Interactions Are Highly Sensitive to Environmental Context

**DOI:** 10.1101/079251

**Authors:** Angela Liu, Anne Archer, Matthew B. Biggs, Jason A. Papin

**Affiliations:** Department of Biomedical Engineering, University of Virginia, Charlottesville, Virginia; Department of Biology, University of Virginia, Charlottesville, Virginia

## Abstract

Microbial interactions are ubiquitous in nature, and equally as relevant to human wellbeing as the identities of the interacting microbes. However, microbial interactions are difficult to measure and characterize. Furthermore, there is growing evidence that they are not fixed, but dependent on environmental context. We present a novel workflow for inferring microbial interactions that integrates semi-automated image analysis with a colony stamping mechanism, with the overall effect of improving throughput and reproducibility of colony interaction assays. We apply our approach to infer interactions among bacterial species associated with the normal lung microbiome, and how those interactions are altered by the presence of benzo[a]pyrene, a carcinogenic compound found in cigarettes. We found that the presence of this single compound changed the interaction network, demonstrating that microbial interactions are indeed highly dynamic and responsive to environmental context.

## Introduction

Microbes are rarely alone. Microbial communities are nature’s workhorses, from degrading tree litter in the forest to degrading cheese burgers in the colon [1,2]. Complex communities of bacteria and fungi surround the roots of plants and colonize the surfaces of our teeth [3,4]. Interactions between microorganisms within these communities can entirely determine the overall interaction of the community with the environment. A potent example is the influence of *Clostridium scindens* on *Clostridium difficile* [5]. *C. difficile* is an intestinal pathogen that can reside indefinitely in the intestinal tract of healthy humans alongside hundreds of other species. However, after antibiotic treatment, *C. difficile* often outcompetes its neighbors and produces toxins, causing the host to experience intense diarrhea, fever and weight loss. Interestingly, the addition of a single species—*C. scindens*—to the intestinal community can prevent *C. difficile* overgrowth and the associated negative symptoms [5]. A single interaction means the difference between health and disease. Similarly, disease severity can be influenced by microbial interactions. For example, *Burkholderia cenocepacia* increases mortality among cystic fibrosis patients with *Pseudomonas aeruginosa*-associated lung infections, illustrating that interactions among microbes are just as relevant to human health as the identities of the species with which we interact [6].

However, it is difficult to measure and characterize microbial interactions [7]. Interactions have many underlying causes including direct interactions through signaling or toxin molecules, competition and cross-feeding, or the result of one species actively changing environmental factors such as pH [8]. Interactions are particularly difficult to measure in complex, multi-species communities. Recent efforts to elucidate the network of microbial interactions relied on time-series measurements of metagenomic sequence data, inferring interaction information from microbial abundance dynamics [9,10]. While useful, these approaches often assume that the nature of microbial interactions is fixed, an assumption which can mask important biological insights. Returning to the interaction between *C. scindens* and *C. difficile*, it was shown that *C. scindens* prevents *C. difficile* overgrowth by producing secondary bile acids, which is only possible in the presence of primary bile acids [5]. Clearly, in at least some cases, the chemical and nutritional context of the environment determines the interactions which are possible. The question of “how fixed are microbial interactions?” is still open, and answering this question requires innovation in the ways we measure microbial interactions, and requires many more observations of microbial communities in many different contexts.

We present a novel screening approach to quantify microbial interactions *in vitro*, enabling us to measure how those interactions change as a function of the environment. As a test case, we chose to examine the influence of benzo[a]pyrene (BaP) on the interactions between a subset of “core” lung bacterial species [11,12]. BaP is a polycyclic aromatic hydrocarbon that is found in cigarette smoke which can interfere with DNA replication [11]. We quantified the pairwise interactions between bacterial species in control and BaP media conditions and found that at least one interaction was completely altered from growth-promoting to growth-inhibiting, while many other interactions changed more subtly in sign or magnitude. This proof-of-principle demonstration highlights the utility of our new framework and the fact that microbial interactions are highly dynamic and responsive to the environment. Improved tools will lead to greater awareness and understanding of microbial interactions, and to an increased ability to target them therapeutically.

## Materials and Methods

### Species Selection

Previous research suggests the existence of a core microbiome in the human lung [12]. We selected bacterial species which are associated with the respiratory tract: *P. aeruginosa* PA01, *P. aeruginosa* PA14, *Haemophilus influenza* type B ATCC 10211, *Haemophilus parainfluenzae* ATCC 7901, and *Staphylococcus aureus* ATCC 29213.

### Media and Culture Protocol

All five species were cultured in brain-heart infusion (BHI) medium (BD) supplemented with L-histidine (0.01 g/L) (Sigma), hemin (0.01 g/L) (Sigma) and β-NAD (0.01 g/L) (Sigma) [13]. For the agar plates, we added granulated agar (BD) at 1.2% by weight. We prepared a stock solution of BaP (Sigma) dissolved in DMSO at 10 mg/mL and filter sterilized this solution (0.2 µm pore size). We added 250 mL of this solution into 1L of supplemented BHI for BaP conditions.

On day 0, we made the agar plates and the liquid medium. We allowed liquid cultures to grow for 24 hours in a shaking incubator at 37°C and 5% CO2. On day 1, we collected OD600 measurements for each of the liquid cultures, diluted the liquid cultures to equal OD600 with fresh medium, and evenly spread 7 mL on agar plates to create a lawn of each species which grew for 24 hours. On day 2, each species was stamped onto fresh 6-well agar plates using a custom stamping mechanism which ensured equal initial spacing and colony size (Fig 1). Each species was grown alone and in pairwise combinations with the other four species. The stamped 6-well plates were placed in the incubator for 24 hours before imaging (Fig 1).

**Fig 1:**
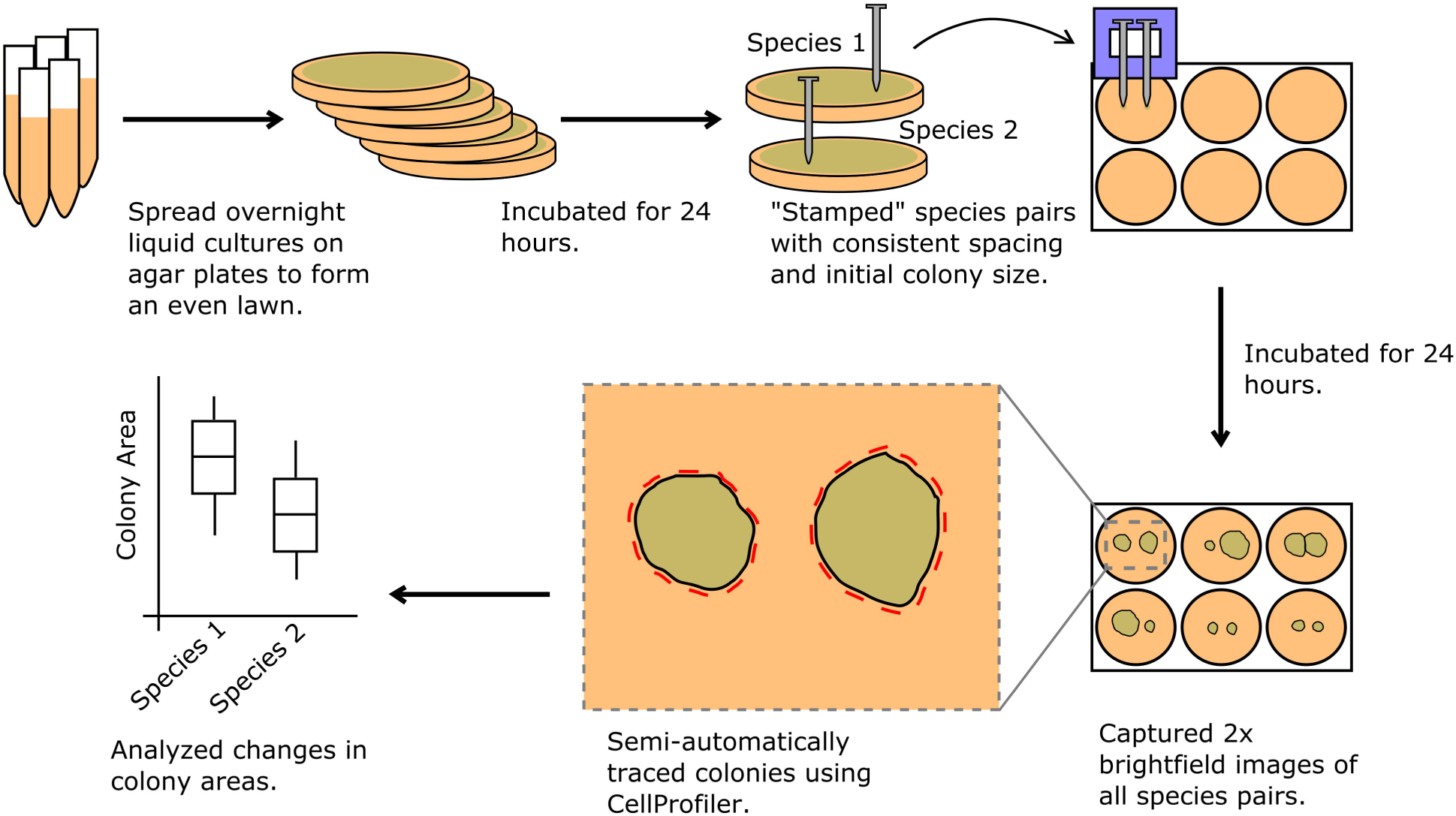
Workflow description. All five species were grown in overnight liquid cultures. Cultures were diluted with fresh media, spread evenly over agar plates and incubated for 24 hours to grow into lawns. We devised a stamping mechanism to maintain consistent spacing and sizes of initial colonies. Nails were used to lift cells from the lawns. The nails were placed into slots in a 3D-printed stamping mechanism and cells were carefully “stamped” onto fresh BHI agar poured in 6-well plates. In the control condition, all five species were grown alone, and all pairwise combinations were grown together (six replicates were performed of each condition). For the BaP condition, the procedure was identical except that BaP was added to the growth media. The stamped colonies were incubated for 24 hours and the resulting colonies were imaged at 2x magnification. The images were automatically segmented and colony areas calculated using image analysis software (see Materials and Methods).

### Stamping Mechanism

In measuring bacterial areas as one of our final metrics, it was necessary to ensure that the initial bacterial colonies were stamped at a consistent size and spacing. We selected a starting spot size of 0.5 mm diameter, which were placed 3.5 mm apart (from center to center). To achieve these specifications, we used metal nails to pick up the bacteria from the lawn and a 3D-printed mechanism to stamp the bacteria onto the plates. The metal nails had uniform tip size and were non-porous, which improved initial inoculation consistency. The stamping mechanism had two fixed slots, each sized to allow the nails to slide in. These slots ensured that the two nails were the same distance apart for each stamping. The nails were autoclaved and the stamping mechanism was routinely disinfected with CaviCide (Metrex).

### Imaging Bacteria

Our images were captured by an EVOS microscope (ThermoFisher Scientific) at 2x magnification and using the brightfield mode (Fig 1). We collected images for six biological replicates of each condition (Fig 2). In order to convert colony areas from pixels to mm^2^, additional images were taken of size standards.

**Fig 2.**
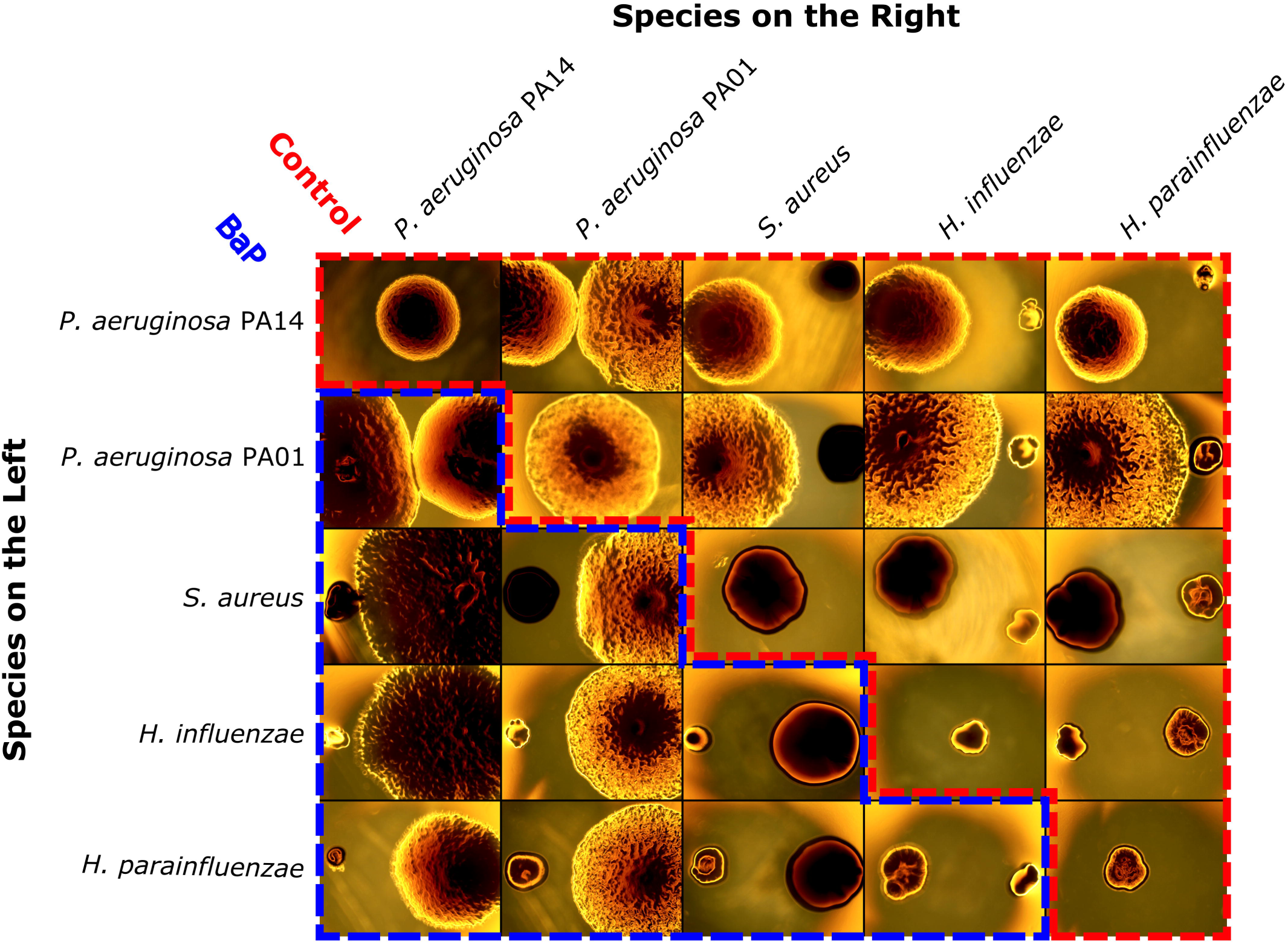
Representative images of bacterial colonies. After being stamped and grown for 24 hours, we imaged all the experimental conditions (six replicates each of five species grown alone and in pairs, in control and BaP conditions). Here we show representative images of the five species grown alone in control media (along the diagonal), the pairwise combinations in control media (upper triangular portion outlined by red, dashed line), and pairwise combinations in BaP media (lower triangular portion outlines by blue, dashed line). The row indicates the species grown on the left and the column indicates the species grown on the right. Interactions were determined by comparing colony area after growth alone to colony area after growth in the presence of another species.

### Data Analysis

Images were analyzed in CellProfiler, an open-source software package [14]. While bacterial colonies were traced automatically in CellProfiler, we manually checked each image to ensure that edge traces were accurate. We collected measurements of colony area (in pixels), shape and location. Colony areas were converted from pixels to cm^2^ based on the images of paper standards (Fig 3). In order to infer microbial interactions, colony areas in paired growth conditions were compared to colony growth alone using a two-sided Wilcoxon signed-rank test in R [15]. The p-values produced were adjusted for multiple testing using the false discovery rate (FDR) method. Similarly, we compared the interactions from the control condition to the condition with BaP exposure in order to identify cases where the nature of an interaction changed. To accomplish this, we converted colony areas to area fold changes by dividing the observations by the mean area when the species was grown alone. The observed fold changes in control versus BaP conditions were compared using a two-sided Wilcoxon signed-rank test and p-values were adjusted for multiple testing using the FDR method.

All of our images and analysis scripts are available in our online repository: https://github.com/mbi2gs/BaP_microbialInteractions

**Fig 3.**
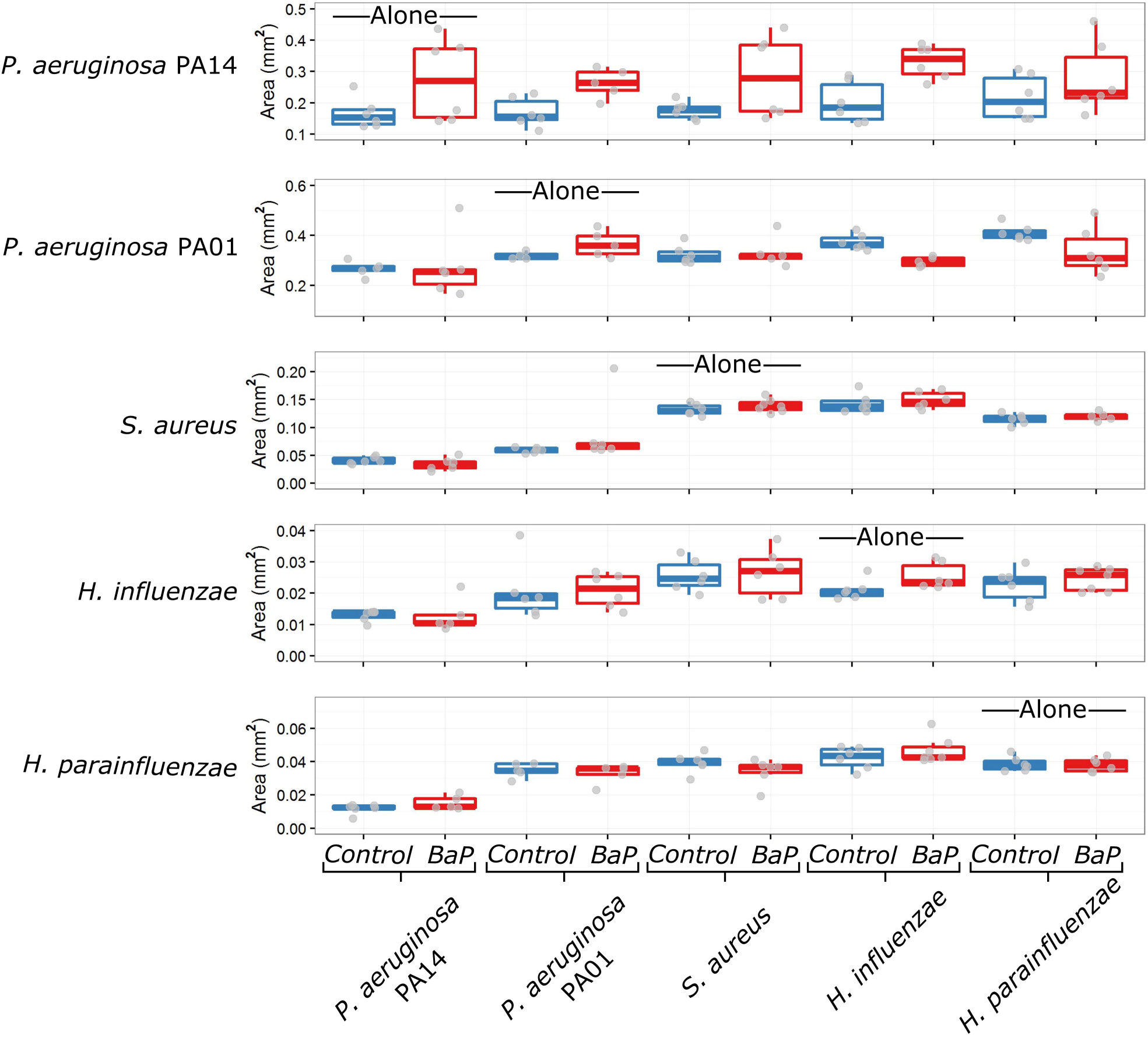
Final colony areas across all conditions. Colony areas were obtained from the images using semi-automated image analysis in CellProfiler. Areas were converted from pixels to mm^2^ based on images of paper standards. Control conditions are shown in blue while BaP conditions are shown in red. Individual data points are shown in gray (note that some outliers are outside the axis range of these plots). The colony areas correspond to the species labels on the left, while the labels along the bottom indicate the paired species. Species interactions are inferred by comparing pairwise growth with growth alone. For example, notice that when grown next to *P. aeruginosa* PA14, all the other species reach smaller colony areas than when grown alone, suggesting that *P. aeruginosa* PA14 negatively interacts with all four other species.

## Results

### Interspecies Interactions Observed in the Control Condition

Final colony areas of four of the five species were significantly altered by at least one interaction in the control media condition (p-value < 0.05 by two-sided Wilcoxon signed-rank test corrected by FDR). We observed that both of the *P. aeruginosa* species grew more when paired with *H. parainfluenzae* or *H. influenzae* in control media, and the increase of growth was statistically significant in the case of *P. aeruginosa* PA01 (Fig 4A). *S. aureus* grew less when it was paired with both of the *P. aeruginosa* species. *P. aeruginosa* PA01, *S. aureus*, *H. parainfluenzae* and *H. influenzae* all grew less when paired with *P. aeruginosa* PA14.

**Fig 4.**
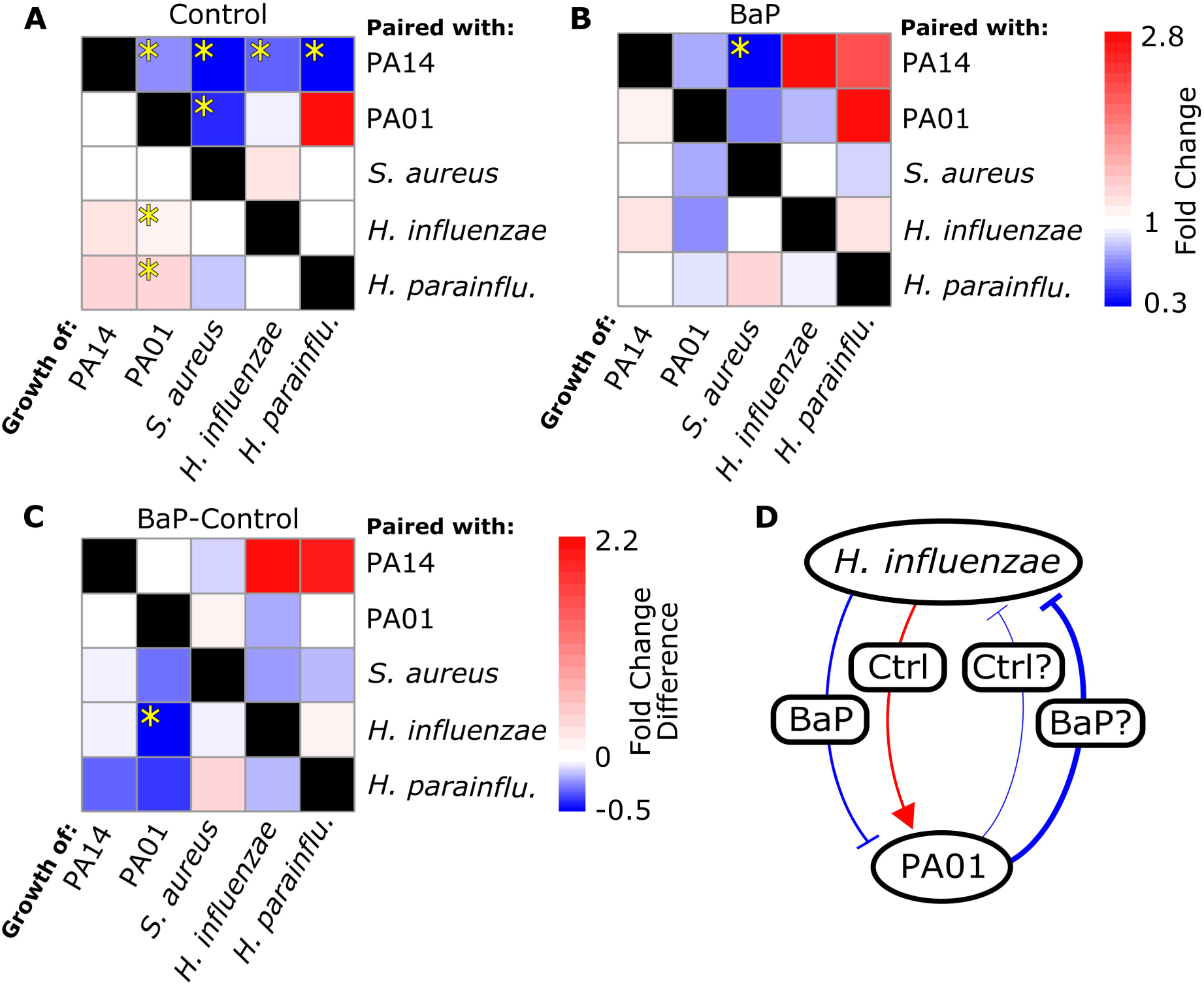
Interactions between species are context-dependent. A–B. Heat maps indicate the fold change in colony area during pairwise growth with respect to growth alone. Fold changes in colony area are shown for the species in the columns. The species along the rows are those which influenced the growth of the species in the columns. Statistically significant changes in colony areas are indicated by yellow stars (p-value < 0.05 by two-sided Wilcoxon signed-rank test corrected by FDR). **A**. Fold changes in colony area in control media. *S. aureus* grows significantly less when grown next to *P. aeruginosa* PA01, and *P. aeruginosa* PA01 grows significantly less when paired with *P. aeruginosa* PA14. **B**. Fold changes in colony area in BaP media. Only one interaction is statistically significant in this context, and it is an interaction that was also observed in the control condition. **C**. Panel A subtracted from panel B. The difference in fold change is an indication of how different the interactions are in the two conditions. Statistical significance was determined by two-sided Wilcoxon signed-rank test on the fold change data for each interaction, and then FDR corrected. Only one interaction was significantly changed between the two conditions. *P. aeruginosa* PA01 grew more when paired with *H. influenzae* in control media, but grew less in BaP media. Some interactions were essentially identical between the two conditions (e.g. *P. aeruginosa* PA01 grows less when paired with *P. aeruginosa* PA14, regardless of the presence of BaP), and others were changed, if not in a statistically significant way (e.g. *P. aeruginosa* PA01 grows more when paired with *H. parainfluenzae* in control media, but less in BaP media). **D**. An illustration of the BaP-dependent interactions between *H. influenzae* and *P. aeruginosa* PA01. We put question marks beside the interactions which are observed but not statistically significant. Note that BaP causes *H. influenzae* to switch from a positive to a negative influence on *P. aeruginosa* PA01.

### BaP Alters Interspecies Interactions

We did not observe statistically significant changes in bacterial colony growth when grown alone in the presence of BaP after correcting for multiple testing, although the increase in *H. influenzae* colony size was significant before FDR correction (Fig 3). In terms of interactions, we observed that *S. aureus* grew significantly less when paired with *P. aeruginosa* PA14 in the BaP media condition (Fig 4B; p-value < 0.05 by two-sided Wilcoxon signed-rank test corrected by FDR). This same interaction (*P. aeruginosa* PA14 inhibition of *S. aureus*) was also observed to be statistically significant in the control condition (Fig 4A). Some interactions from the control condition, while no longer statistically significant in the BaP condition, did retain the same trends from the control condition. *P. aeruginosa* PA01 still grew less when paired with *P. aeruginosa* PA14 and *S. aureus* growth was inhibited when paired with *P. aeruginosa* PA01 (Fig 4B). The trend of other interactions from the control condition seemed to change in the BaP condition (Fig 4C). The only statistically significant change was the interaction between *H. influenzae* and *P. aeruginosa* PA01. In the control condition, *P. aeruginosa* PA01 grew more when paired with *H. influenzae*, while in the BaP condition, *P. aeruginosa* PA01 grew less when paired with *H. influenzae* (Fig 4C). A similar trend can be seen in the interaction between *H. parainfluenzae* and *P. aeruginosa* PA01, although it is not statistically significant after multiple testing correction.

## Discussion

We devised an *in vitro* experimental framework that allows for the semi-automated assessment of pairwise interactions between microbes grown *in vitro*. We demonstrate our framework on a set of five representative lung bacterial species. We identified several growth-altering interactions in both control and BaP conditions, and significantly, we saw that the nature of at least one observed interaction was significantly different between the two conditions, indicating that interspecies interactions are altered by the presence of BaP.

Our pipeline includes a novel stamping mechanism for standardizing colony sizes, combined with quantitative image analysis (Fig 1). We incorporated the open-source image analysis software CellProfiler into our workflow to increase throughput and quantify interactions [14]. Our methodology allows for automated, quantitative measurements of colony attributes whereas previous research using similar techniques has relied on manual observations [16]. For this work, we chose to focus on colony area as an indicator of interaction strength. Colony area is a function of both growth rate and duration. We assume that a bacterial colony will grow more slowly and/or stop growing earlier when the cells are under stress (due to some negative interaction). Similarly, we assume that a colony will grow faster and/or longer if it is participating in a beneficial interaction. Our workflow does not give much insight into the mechanism of the interaction except perhaps to require that interactions act over a distance, since generally the colonies are not physically interacting. Potential diffusion-based causes of interactions include (but are not limited to) metabolic competition, cross-feeding, signaling, or toxin-mediated interactions.

In this study we observed many interspecies interactions, some of which have never been reported in the literature. In both control and BaP conditions neighboring species grew less when paired with *P. aeruginosa* PA14 (and to a lesser extent, PA01; Fig 3). One contributing factor is that *P. aeruginosa* PA14 and PA01 colonies were enormous (Fig 2), suggesting that these two species utilized nutritional resources more rapidly than neighboring species could access them. It is also known that many species of *P. aeruginosa* excrete molecules that broadly inhibit growth among neighboring species, including *S. aureus* and *Haemophilus* species [16–18]. We also observed that both *H. influenzae* and *H. parainfluenzae* promote growth of *P. aeruginosa* PA01 (and to a lesser extend PA14), an interaction which to our knowledge has never been reported before (Fig 4A), highlighting the utility of our approach for identifying relevant and novel interspecies interactions.

An interesting outcome was the sensitivity of the observed interactions to the presence of BaP. In the control condition, *P. aeruginosa* PA01 growth was enhanced when paired with *H. influenzae*, while in the presence of BaP, *P. aeruginosa* PA01 growth was inhibited when paired with *H. influenzae*, completely reversing the nature of the interaction (Fig 4D). While this reversed interaction was the most dramatic observation, many interactions changed in nature or intensity to some extent, if not statistically significantly (Fig 4C). It is remarkable that the addition of a single chemical could so drastically alter elements of the interaction network, which has important implications in the context of human health where the effect of pharmaceuticals on the human microbiota are rarely understood [19]. There is growing evidence that microbial interactions are not fixed, but are context-dependent and in continuous feedback with the local chemical backdrop [16,20,21]. Computationally, it has been shown that pairs of bacterial species can be made to compete or cooperate based solely on the nutritional context in which they are grown [22]. The context-dependence of bacterial interactions becomes relevant during attempts to predict the dynamics of important microbial communities such as the gut microbiome. Previously, mathematical frameworks for inferring microbial interaction networks have been based on the assumption that interactions are fixed [10,23]. Advances in predicting and engineering community dynamics will come as we increase our understanding of the effect of context on microbial interactions.

Future applications of our workflow can easily utilize metrics other than colony area. Readily-included metrics include colony shape and color. While we did not observe visible phenotypic differences in this particular study, previous research suggests that shape and color can reveal important interactions. For example, pseudomonads can exhibit a “swarming” phenotype which drastically alters colony shape, and *Streptomyces coelicolor* can produce a large variety of pigments which differ depending on the identity of neighboring species [24,25].

The novel pipeline applied here enables faster screening of microbial interactions *in vitro*. Our results derived from representative lung bacteria highlight the fact that microbial interactions are dynamic and responsive to environmental perturbations. Improved tools such as those presented here will lead to greater understanding of such interactions, and to an increased ability to modulate them therapeutically.

## Acknowledgements

We gratefully acknowledge funding for this work through the University of Virginia by a Harrison Undergraduate Research Award to AL and AA. This work was further supported by NIH grant [5R01GM108501] to JP.

